# A flexible repertoire of transcription factor binding sites and diversity threshold determines enhancer activity in embryonic stem cells

**DOI:** 10.1101/2020.04.17.046664

**Authors:** Gurdeep Singh, Shanelle Mullany, Sakthi D Moorthy, Richard Zhang, Tahmid Mehdi, Ruxiao Tian, Alan M Moses, Jennifer A Mitchell

## Abstract

Transcriptional enhancers are critical for development, phenotype evolution and often mutated in disease contexts; however, even in well-studied cell types, the sequence code conferring enhancer activity remains unknown. We found genomic regions with conserved binding of multiple transcription factors in mouse and human embryonic stem cells (ESCs) contain on average 12.6 conserved transcription factor binding sites (TFBS). These TFBS are a diverse repertoire of 70 different sequences representing the binding sites of both known and novel ESC regulators. Remarkably, using a diverse set of TFBS from this repertoire was sufficient to construct short synthetic enhancers with activity comparable to native enhancers. Site directed mutagenesis of conserved TFBS in endogenous enhancers or TFBS deletion from synthetic sequences revealed a requirement for more than ten different TFBS. Furthermore, specific TFBS, including the OCT4:SOX2 co-motif, are dispensable, despite co-binding the OCT4, SOX2 and NANOG master regulators of pluripotency. These findings reveal a TFBS diversity threshold overrides the need for optimized regulatory grammar and individual TFBS that bind specific master regulators.

## INTRODUCTION

Sequence variation in transcriptional enhancers modulates inter and intra-species phenotypic divergence due to changes in gene expression (Kvon et al., 2016; McLean et al., 2011; Prescott et al., 2015). These non-coding regulatory sequences are required to regulate tissue specific gene expression during normal development and genome-wide association studies (GWAS) reveal that disease-associated SNPs (single nucleotide polymorphisms) are more often found in non-coding regions of the genome displaying chromatin features associated with transcriptional enhancers (Bernstein et al., 2012; Maurano et al., 2012; Visel et al., 2009a). It is known that transcription factor binding sites (TFBS) are required for enhancer function, and that transcription factors, modulate enhancer activity in a cell type specific manner; however, the precise sequence code conferring enhancer activity in each cell type remains unknown.

Extreme sequence conservation between mouse and human non-coding regions identifies active enhancers during development (Visel et al., 2008) but most mammalian enhancers do not display high sequence conservation. Enhancers active in specific cell types can be identified using chromatin features; for example, accessible chromatin, histone modifications, transcription factor or co-activator binding (Chen et al., 2012, 2008; Ernst and Kellis, 2012; Libbrecht et al., 2019; Visel et al., 2009b). Histone H3 K27 acetylation (H3K27ac) and H3 K4 monomethylation (H3K4me1) are enriched at enhancer regions, with increased H3K27ac signal associated with more active enhancers (Creyghton et al., 2010; Moorthy et al., 2017; Rada-Iglesias et al., 2011). However, predictions based on these chromatin features alone include ~75% false positives (Barakat et al., 2018; Corces et al., 2018; Farnham, 2012; Kwasnieski et al., 2014; Visel et al., 2008, 2013) and analysis of the identified enhancers has yet to reveal a DNA sequence code that determines enhancer function. Comparative chromatin analysis of liver from 20 mammals revealed that regions with conserved H3K27ac in multiple species (Villar et al., 2015) can be exploited to identify conserved enhancer sequence properties (Chen et al., 2018).

Regions bound by multiple transcription factors, display increased regulatory activity by enhancer deletion, transgenic reporter assays and massively parallel reporter assays (MPRAs), compared to regions bound by single transcription factors (Moorthy et al., 2017; Vanhille et al., 2015; Zinzen et al., 2009). In addition, regulatory activity is higher for synthetic heterotypic sequences constructed from four different TFBS compared to homotypic sequences containing single repeated TFBS (Fiore and Cohen, 2016; Smith et al., 2013), however, the activity of these synthetic sequences has not been benchmarked against native enhancers. Combinatorial regulation by transcription factors is important for enhancer function; however, only 37% of genomic regions containing TFBS for 6 different regulatory transcription factors display activity (Lloret-Fernández et al., 2018). Studies of specific native enhancers and MPRAs reveal orientation and spacing of TFBS pairs, often referred to as a regulatory grammar, affects enhancer activity (Farley et al., 2016; Fiore and Cohen, 2016; Thanos and Maniatis, 1995). This regulatory grammar, in combination with TFBS clustering, however, does not predict which regions exhibit enhancer activity in the genome indicating we have an incomplete understanding of the regulatory code (King et al., 2020; Lusk and Eisen, 2010; Smith et al., 2013).

As regions bound by the same transcription factor in multiple species are more likely to have regulatory activity (Ballester et al., 2014) we investigated regions with conserved binding of multiple transcription factors in mouse and human embryonic stem cells (ESCs) to decipher the pluripotency regulatory code. This analysis revealed a large repertoire of 70 different TFBS contribute to enhancer activity in pluripotent cells. Within individual enhancers more than ten different TFBS are required for robust enhancer activity, indicating TFBS sequence diversity is a required feature of mammalian enhancers.

## RESULTS

### Comparative epigenomics reveals that conserved enhancers in pluripotent stem cells contain a conserved regulatory code

To decipher the regulatory code in pluripotent cells, we investigated regions bound by multiple transcription factors in mouse and human ESCs (* in Figure 1A), as these regions are more likely to show enhancer activity (Ballester et al., 2014). To identify these regions, 1 kb segments bound by at least one transcription factor in mouse were clustered based on active enhancer features: H3K27ac (Creyghton et al., 2010; Rada-Iglesias et al., 2011) and transcription factor binding in mouse and human ESCs (Figure 1B, Table S1). The resulting clusters included regions with conserved high enhancer features (CHEF), conserved medium enhancer features (CMEF), mouse-specific high enhancer features (msHEF), and several conserved low enhancer feature (CLEF) clusters, most of which were bound by only one transcription factor and displayed low H3K27ac in both species. The CHEF and msHEF clusters, containing the greatest enrichment for active enhancers (Moorthy et al., 2017; Murtha et al., 2014; Schnetz et al., 2010; Zhou et al., 2014), and increased association of the EP300 co-activator (Figure 1C), make up only 17% of the total mouse transcription factor bound regions in accordance with a recent MPRA revealing only 17-26% of transcription factor bound regions possess enhancer activity (Barakat et al., 2018). By starting with transcription factor bound regions in mouse we could not identify human-specific high enhancer feature (hsHEF) regions; however, clustering transcription factor bound regions in human ESCs identified hsHEF regions, and 86.5% of the same CHEF regions identified in mouse (Figure S1A).

**Figure 1:**
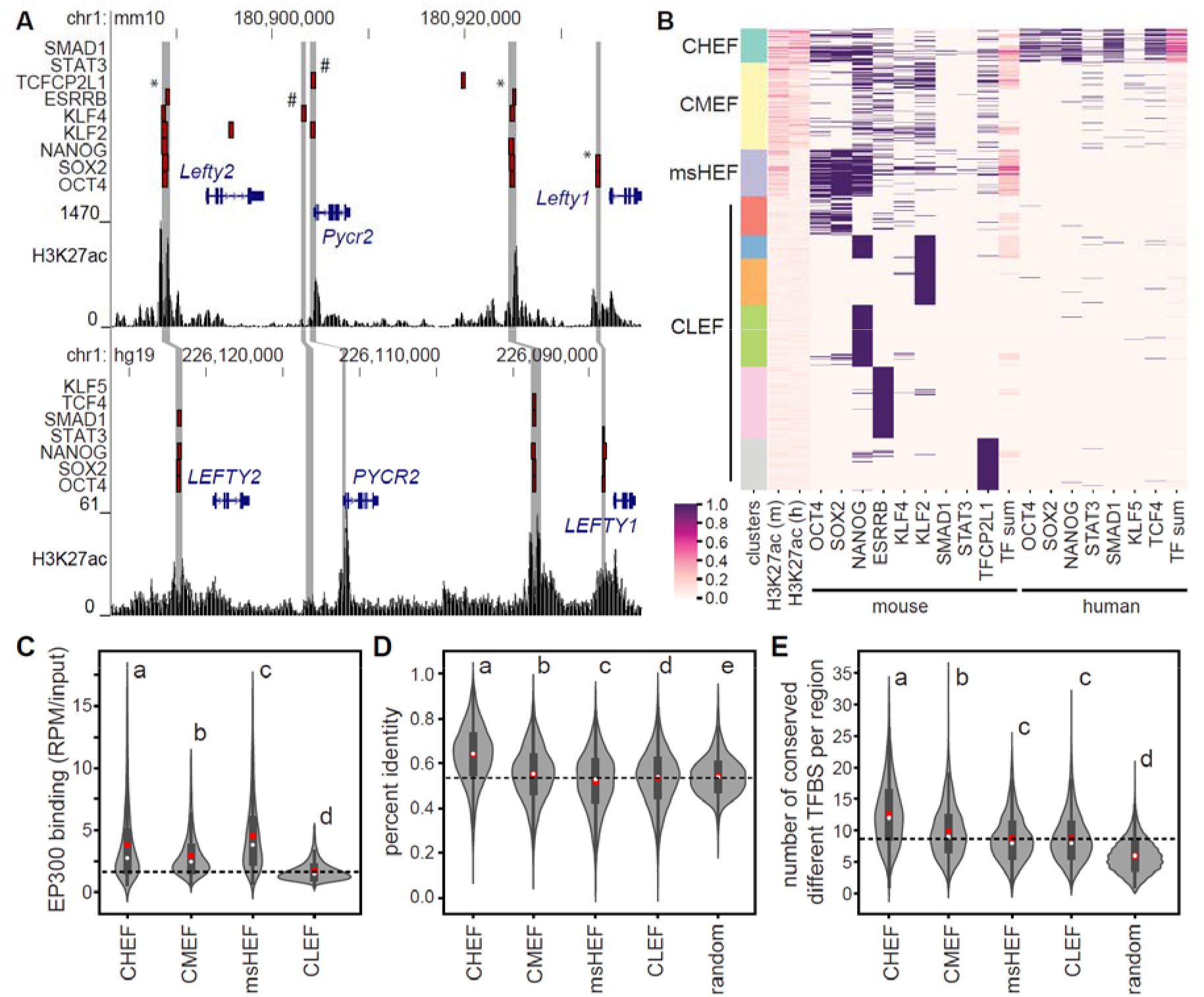
Conserved high enhancer feature (CHEF) regions contain a conserved regulatory code. A) The *Lefty1*/*Lefty2* locus in the mouse (top) and human (bottom) genomes. Transcription factor bound regions from ChIP-seq (red bars), mouse and human syntenic regions (grey bars), and H3K27ac ChIP-seq data are displayed on the mm10 and hg19 assemblies of the University of California at Santa Cruz (UCSC) Genome Browser. * indicates regions with conserved binding of multiple transcription factors in mouse and human, # indicates regions with binding only in mouse ESCs. B) Clustering of transcription factor bound regions in mouse ESCs, using H3K27ac, transcription factor binding, and the number of transcription factors bound in a region (TF sum) at associated mouse and human regions. In C-E groups determined by one-way ANOVA to be significantly different (P<0.05) are labelled with different letters, dashed line indicates the average for the CLEF cluster. C) CHEF, CMEF and msHEF regions display significantly increased EP300 binding compared to CLEF regions. D) CHEF regions display the highest percent identity between mouse and human compared to other clusters and random non-transcription factor bound regions. E) CHEF regions contain an increased number of conserved different TFBS compared to other clusters and random non-transcription factor bound regions.

We next identified features that distinguished CHEF sequences from other genomic regions. CHEF sequences had increased DNA sequence conservation compared to other clusters, but could not be identified based on conservation alone (Figure 1D). As MPRA revealed that TFBS conservation, rather than overall conservation is a determinant of enhancer activity (Kheradpour et al., 2013), we analyzed matches to known transcription factor binding motifs. CHEF mouse-human TFBS conservation by comparison to scrambled TFBS indicated several TFBS are significantly conserved (P<2.2×10^−16^, Wilcoxon rank-sum test, Figure S1B/C). We tested this further by evaluating TFBS conservation in CHEF sequences across six species using MotEvo (Arnold et al., 2012). As TFBS diversity is important for activity of synthetic regulatory sequences (Fiore and Cohen, 2016; Smith et al., 2013), we evaluated the number of different conserved TFBS, corresponding to ESC expressed transcription factors based on RNA-seq, in each of the sequences. From this set of 349 TFBS, we found CHEF sequences contain on average 12.6 conserved and different TFBS, an increased number compared to all other groups (Figure 1E). Even after removal of TFBS associated with transcription factors for which binding data was used to identify CHEF regions, CHEF sequences contain an increased number of conserved and different TFBS compared to all other groups, indicating there are additional regulatory transcription factors important in pluripotent cells (Figure S1D).

### Conserved enhancers contain a large repertoire of TFBS and limited conserved regulatory grammar

To determine the importance of specific TFBS, LASSO (least absolute shrinkage and selection operator)(Tibshirani, 1996) logistic regression was used to identify the TFBS enriched and conserved in CHEF sequences compared to the NANOG bound low enhancer feature cluster (chosen because NANOG lacks a specific TFBS in JASPAR, see Methods). This analysis identified 125 enriched and conserved TFBS, which based on high correlation between PWM (position weight matrices) could be reduced to 70 unique TFBS (see methods, Figure 2A, S2A and Table S2). These include TFBS bound by well characterized pluripotency associated transcription factors (for example: OCT4:SOX2, KLF4, ESRRB) and those corresponding to ChIP-seq data used for clustering (Figure 1B), as well as several novel TFBS. To test enhancer regions for enriched binding of transcription factors corresponding to novel CHEF-enriched TFBS, we identified ChIP-seq data for PRDM14, ZFX and E2F1 in ESCs. These data were not included in the initial clustering as these transcription factors are not considered core pluripotency regulators. PRDM14, ZFX and E2F1 all display enriched binding in the enhancer positive (CHEF, CMEF and msHEF) regions compared to the negative CLEF regions (P<0.0001, hypergeometric test, Figure 2B, S2B). In the case of MTF2, a known repressor (Zhang et al., 2011b), the TFBS was depleted from conserved enhancer regions. Consistent with this finding, ChIP-seq peaks for MTF2 are significantly depleted from CHEF regions compared to CLEF regions (P<0.0001, hypergeometric test, Figure 2B). To evaluate conserved regulatory grammar in CHEF regions we investigated orientation bias for pairs of TFBS from the 70 unique TFBS which were enriched in CHEF regions (Table S3). We determined that only 7.9% (191/2415) of enriched TFBS pairs displayed orientation bias in CHEF regions. These results indicate that CHEF regions are enriched with several conserved and unique TFBS but that TFBS pairs generally do not display conserved relative orientation.

**Figure 2:**
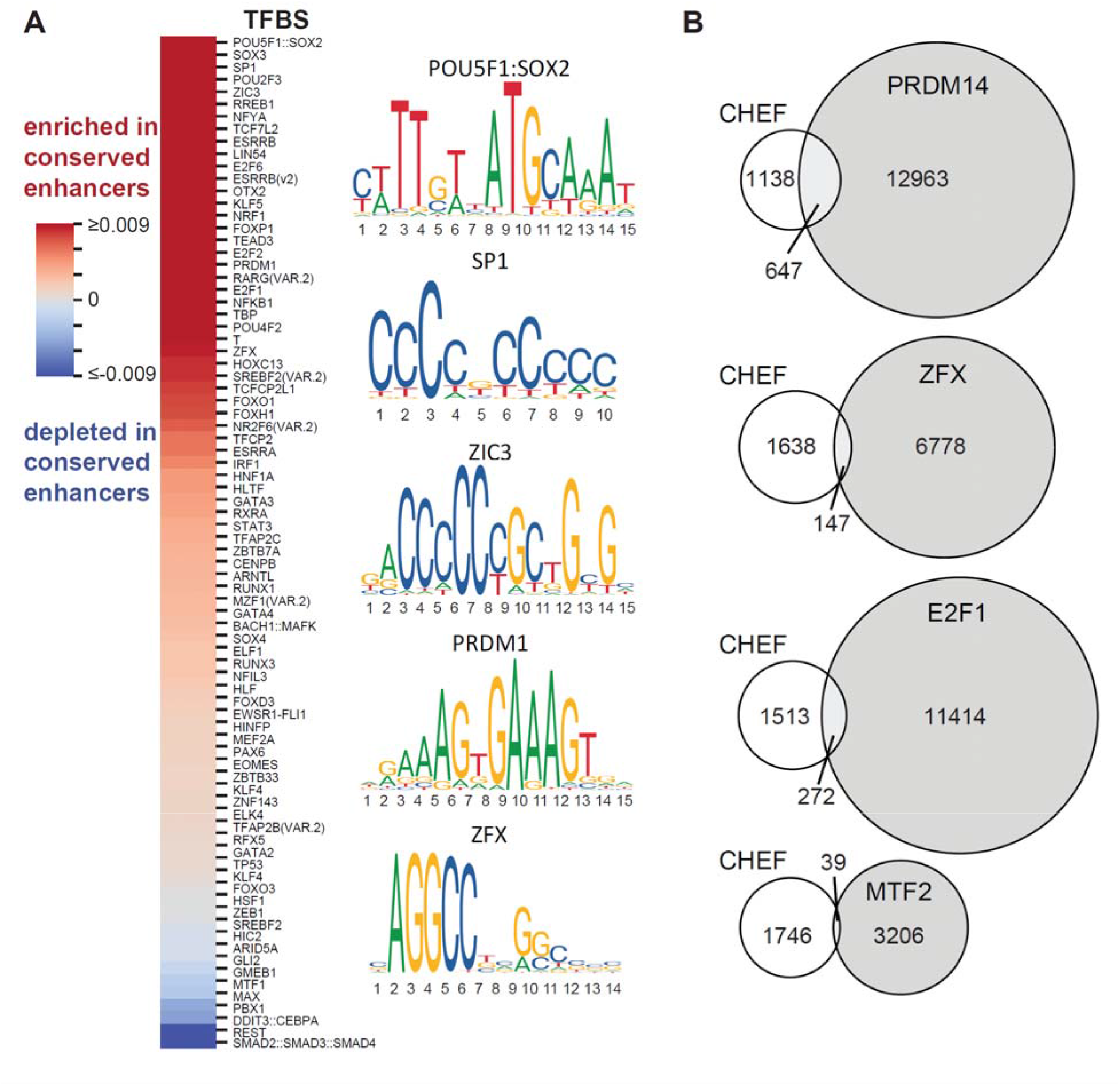
A large repertoire of transcription factor binding sequences contribute to enhancer activity. A) Heatmap indicating the TFBS enriched (red) or depleted (blue) in conserved high enhancer feature regions compared to the NANOG bound negative regions. B) Overlap between conserved high enhancer feature (CHEF) regions and ChIP-seq peaks for transcription factors predicted to bind these regions based on TFBS enrichment (PRDM14, E2F1, ZFX) or predicted not to bind CHEF regions based on TFBS depletion (MTF2).

### Construction of synthetic enhancers reveals transcription factor binding site diversity is required and sufficient for robust enhancer activity

As CHEF regions with conserved transcription factor binding in mouse and human ESCs contain more than 12 conserved TFBS and little evidence of conserved regulatory grammar we hypothesized that TBFS diversity is sufficient for enhancer activity. If this is the case, we should be able to construct synthetic enhancers based on this model. Synthetic sequences containing consensus TFBS separated by 2bp spacers were constructed for evaluation in reporter assays (Table S4). We first evaluated a construct containing 14 copies of the OCT4:SOX2 TFBS (14OS), as this PWM was the most enriched in CHEF sequences. Somewhat surprisingly we determined that this construct had almost no enhancer activity, compared to the native SOX2 enhancer (Figure 3A). To evaluate the role of TBFS diversity we selected TFBS enriched in CHEF regions and created three synthetic enhancers, each containing a different set of 14 TFBS (Table S4). All three sequences displayed robust enhancer activity, comparable to the native *Sox2* enhancer, and activity significantly higher than the synthetic sequence containing 14 copies of the OCT4:SOX2 co-motif (P<0.001, ANOVA, Figure 3A), supporting a heterotypic enhancer model in which multiple different TFBS are needed for robust enhancer activity and sequences composed entirely of repeated TFBS should not act as enhancers. One of the synthetic sequences (14dTFBS_a) contains the OCT4:SOX2 co-motif; however, this motif is dispensable for enhancer activity as the other two synthetic enhancers (14dTFBS_b and 14dTFBS_c) lacked both OCT4 and SOX2 motifs (Figure 3A). Previous optimization of synthetic sequences, containing KLF4, SOX2, OCT4 and ESRRB TFBS, identified two ordered combinations with the highest activity by MPRA (ksOE and sOKE)(Fiore and Cohen, 2016; King et al., 2020). Compared to our synthetic enhancers containing 14 different TFBS and to the native enhancer, these optimized 4TFBS sequences displayed significantly less activity (P<0.01, ANOVA, Figure 3A). This finding suggests that 4 TFBS is below the threshold for robust enhancer activity even when an optimized regulatory grammar is used. To evaluate the role of spacer sequences in enhancer activity we compared the 2bp CC spacers we used to the longer spacers used in the Fiore and Cohen MPRA [ksOE(long) and sOKE(long)] and determined that spacer length does not significantly affect enhancer activity in this context (Figure 3A, Table S4).

**Figure 3:**
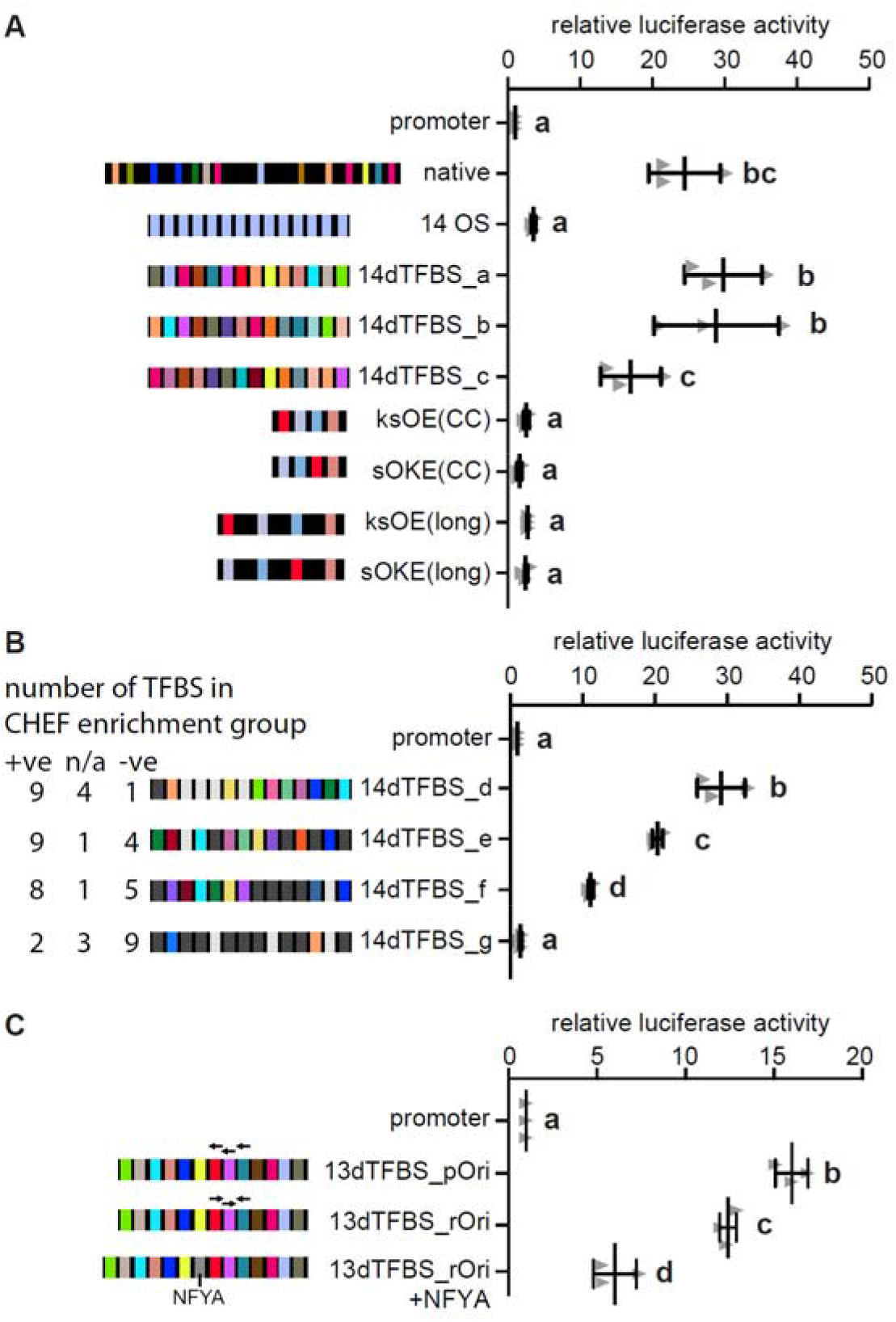
Synthetic sequences reveal transcription factor binding site diversity is required and sufficient for robust enhancer activity. In A-C error bars represent standard deviation, groups determined by one-way ANOVA to be significantly different (P <0.05) are labelled with different letters, n≥3 biological replicates. A) Synthetic enhancers were evaluated in reporter assays and compared to the activity of the *Sox2* enhancer (native). A sequence containing 14 OCT4:SOX2 TFSB (14 OS), 14 different TFBS (14dTFBS_a, _b, _c) from the CHEF-enriched TFBS were evaluated. Optimized 4 TFBS sequences (ksOE, sOKE), with either CC or long spacers between motifs were also evaluated. B) Enhancer activity is reduced when sequences contain fewer CHEF-enriched (+ve) and more CHEF-depleted (−ve) TFBS. The number of TFBS that are neither enriched or depleted are indicated by n/a. C) The effect of motif orientation and repressor binding on enhancer activity. 13dTFBS_pOri contains the preferred orientation, 13dTFBS_rOri contains reversed TFBS. Addition of the repressor NFYA to 13dTFBS_rOri affects but does not abolish enhancer activity.

We hypothesized that a threshold of TFBS corresponding to those enriched in CHEF regions is required for enhancer activity, and furthermore, that TFBS substitution with TFBS depleted from CHEF regions would interfere with activity. We tested this hypothesis with sequences constructed from 14 different TFBS containing a varying number of TFBS enriched, depleted or neutral in CHEF sequences (+ve, −ve, n/a, Figure 3B, Table S2). 14dTFBS_d to _g have an increasing number of TFBS from the CHEF-depleted set (−ve). Enhancer activity was found to be lower in constructs with an increased number of TFBS from CHEF-depleted set (−ve), and fewer TFBS from CHEFenriched set (+ve) (Figure 3B). In fact, 14dTFBS_g had no enhancer activity with 12 TFBS from the CHEF-depleted or neutral set (−ve and n/a) and only 2 TFBS from the CHEF-enriched set (+ve, Figure 3B). All TFBS included in our analysis corresponded to ECS expressed transcription factors, indicating that although there is flexibility in the specific TFBS that contribute to activity, the combination of any 14 TFBS corresponding to ESC expressed transcription factors do not function as an enhancer.

Although we were able to construct active synthetic enhancers by randomly ordering the selected TFBS, we also identified orientation bias for a limited subset of enriched TFBS pairs within CHEF sequences. For example, ZNF263 showed orientation bias with SP1 and KLF4 (P<0.001, Binomial Test, Table S3). To evaluate the effect of orientation a synthetic sequence containing 13 TFBS with the preferred orientation for these TFBS (13dTFBS_pOri) was compared to the reversed orientation (13dTFBS_rOri, Figure 3C). This comparison revealed that orientation has an effect on enhancer activity but is not a requirement as activity was not abolished in the reversed orientation construct. We also observed that FOXJ3 and FOXH1 are more often distant from NFYA in CHEF sequences (Table S3), suggesting pairing the NFYA TFBS with FOXJ3 and FOXH1 could disrupt enhancer activity. This is supported by the observation that NFYA can act as a repressor in combination with specific interaction partners (Peng and Jahroudi, 2002). We placed the NFYA binding sequence adjacent to FOXH1 and FOXJ3 (13dTFBS_rOri+NFYA) and observed a significant reduction in enhancer activity compared to 13dTFBS_rOri (Figure 3C). These data indicate regulatory grammar can affect enhancer activity but has a subtle effect which may fine tune gene expression.

### Additional TFBS confer gain of function to an inactive transcription factor bound region

To test if a diversity of TFBS could confer gain of function to a native inactive transcription factor bound region we evaluated a cluster of regions downstream of *Sall1*. Deletion of the entire cluster affected only *Sall1* transcription determined by RNA-seq (Moorthy et al., 2017), indicating *Sall1* is the only regulated gene for this cluster. Within the cluster (ΔEC) there are three separate transcription factor bound regions designated as multiple transcription factor bound loci (MTL, Figure 4A). Only two of these MTL have enhancer activity in a reporter assay or by CRISPR/Cas9-meidated deletion from the genome (Figure 4B/C). In addition, *Sall1* transcript abundance after deletion of both MTL40 and 28 did not differ from deletion of the entire cluster indicating MTL52 has no independent enhancer activity (Figure S3). Within a core region of MTL52, 6 different transcription factors bind at an overlapping region, whereas the active MTL (40/28) are each bound by 9 transcription factors. We hypothesized MTL52 remained below a required TFBS diversity threshold, but that enhancer activity could be gained by adding a limited number of TFBS. Addition of three TFBS: ESRRB, TFCP2L1 and SMAD3 (Figure 4D, +E+T+S); or TFCP2L1, SMAD3 and E2F1 (Figure 4D, +T+S+E2F1) to the MTL52 core region caused significantly increased enhancer activity, whereas addition of only two TFBS was not sufficient (+T+S, Figure 4D). These data indicate that diversity and abundance of different TFBS are required for enhancer activity, but there is flexibility in TFBS usage.

**Figure 4:**
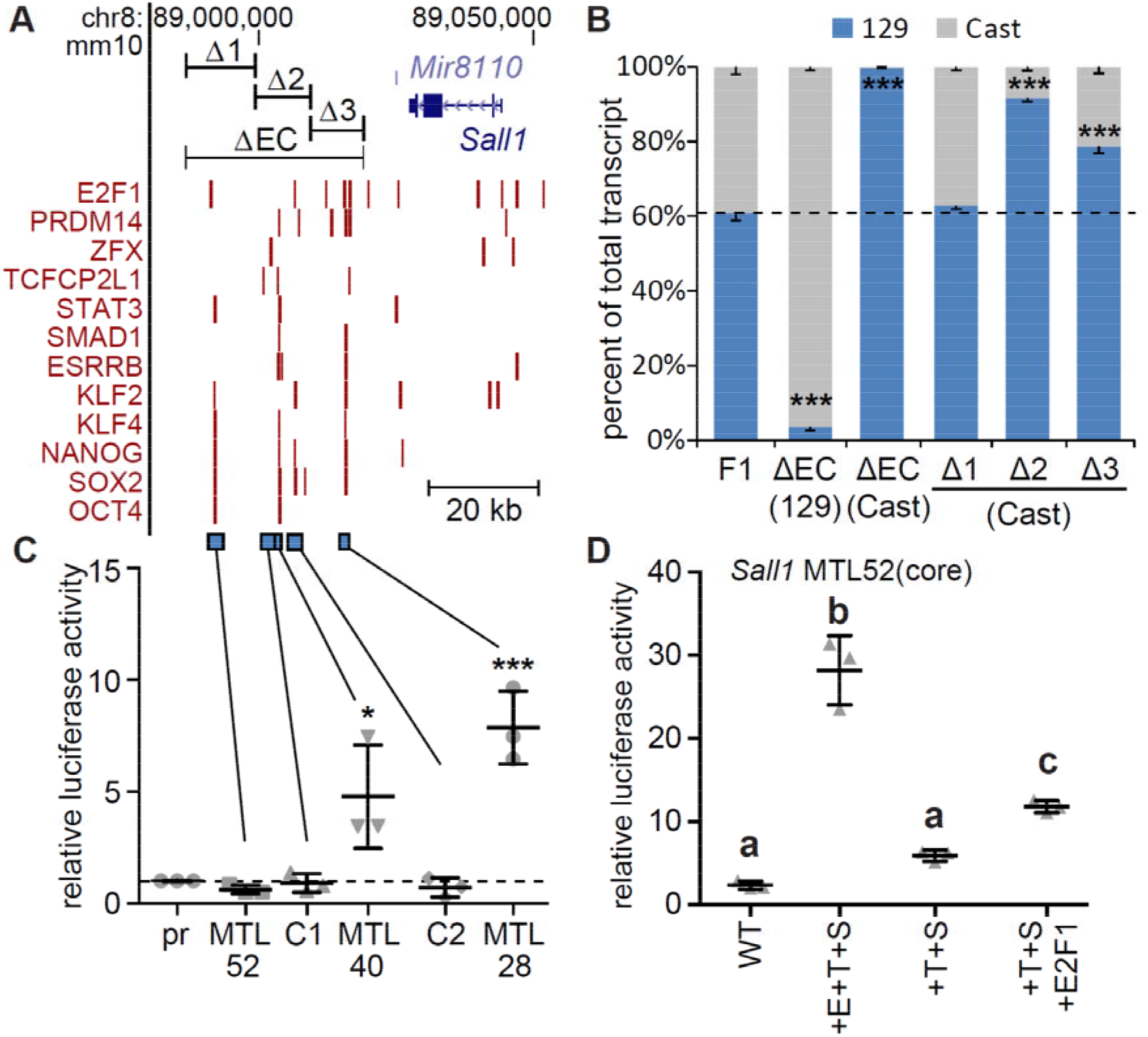
Addition of three different transcription factor binding sequences confers activity to an inactive region bound by six transcription factors. A) Transcription factor bound regions in the *Sall1* locus from ChIP-seq (red bars) are displayed on the mm10 assembly of the University of California at Santa Cruz (UCSC) Genome Browser. At the top CRISPR deleted regions (ΔEC, Δ1, Δ2, Δ3) are displayed. At the bottom regions tested for enhancer activity are displayed (blue bars). B) *Sall1* expression in wild type F1 clones (F1) compared to clones with the indicated deletion. Allele-specific primers detect 129 or Cast RNA in RT-qPCR. Expression for each allele is shown relative to the total. Error bars represent SEM, n≥3 biological replicates. Significant differences from the F1 values are indicated by *** P < 0.001. In C/D error bars represent standard deviation, n≥3 biological replicates. C) Luciferase activity at control (C1, C2) regions and multiple transcription factor bound loci (MTL) 52, 40 and 28 kb downstream of *Sall1*. Significant differences from pr (promoter only) were determined by t-test and are indicated by * P <0.05, *** P<0.001. D) Luciferase activity for wild type (WT) MTL52 core transcription factor bound region, MTL52 core with ESRRB, TCFCP2L1, and SMAD3 (+E+T+S) motifs mutated to the consensus TFBS. From +E+T+S, ESRRB (+T+S) was removed. E2F1 was added to +T+S (+T+S+E2F1). Groups determined by one-way ANOVA to be significantly different (P <0.05) are labelled with different letters.

### A threshold number of TFBS are required for native and synthetic enhancer activity

The number of conserved TFBS in natural enhancers, activity of our synthetic enhancers with a high number (14) of TFBS compared to the few (4) TFBS constructs, and the addition of TFBS to *Sall1* MTL52 all suggest a threshold number of TFBS are required for enhancer activity. To experimentally investigate this threshold more directly, we evaluated synthetic sequences with a decreasing number of different TFBS (Figure 5A, Table S4). Sequential removal of TFBS from 14dTFBS_a revealed that sequences with 14 and 12 different TFBS displayed significant enhancer activity whereas sequences with 5, 7 and 10 different TFBS did not display significant activity. Each of these sequences contains the OCT4:SOX2 TFBS and although this TFBS is the most overrepresented in the CHEF regions, more than 9 additional different TFBS are required for robust activity (12dTFBS, Figure 5A, Table S4) suggesting that more than ten different TFBS are required for robust enhancer activity, even when TFBS bound by master regulatory transcription factors are present in the sequence.

**Figure 5:**
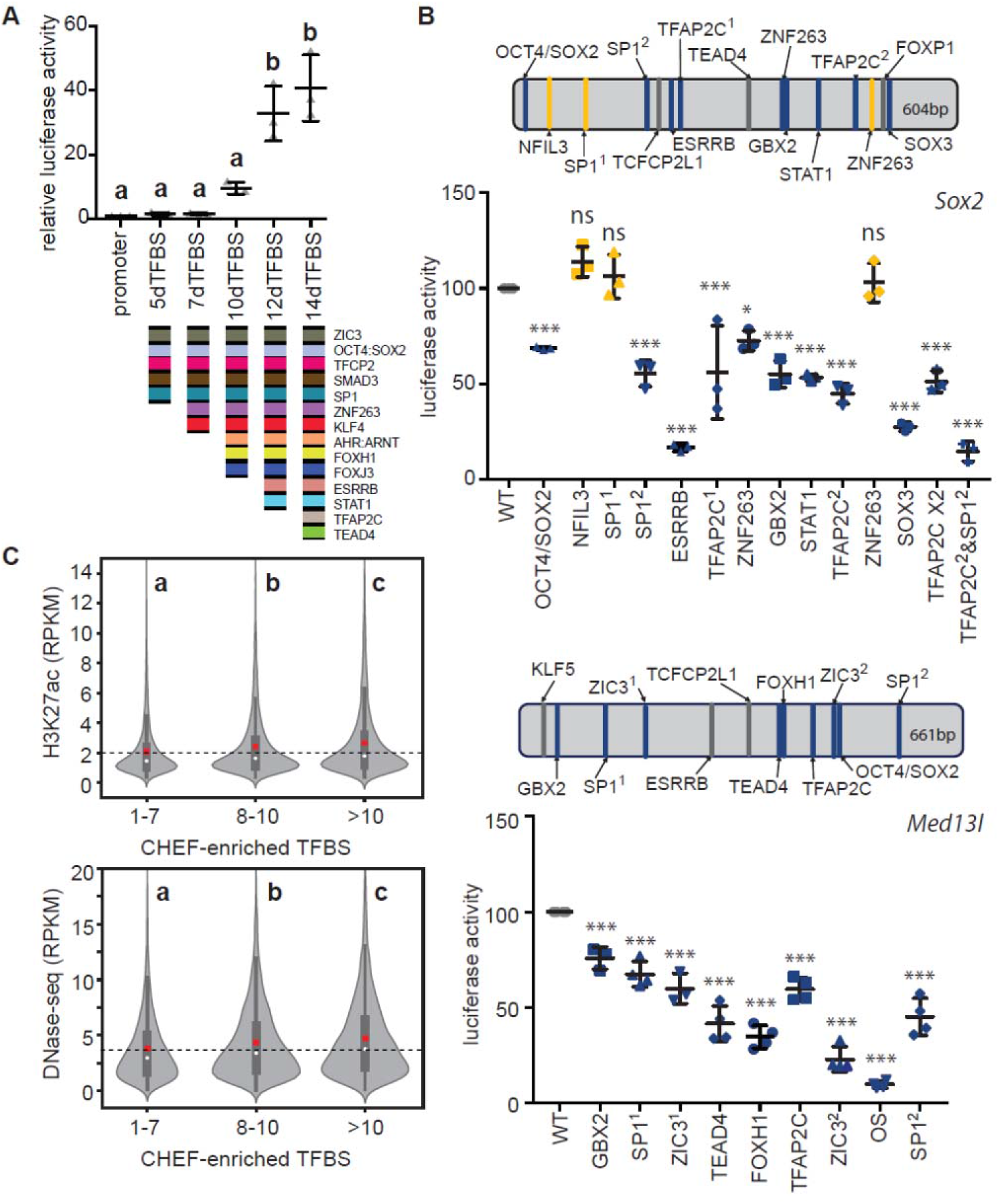
A large repertoire of transcription factor binding sequences and more than ten TFBS are required for enhancer activity. A) Sequential removal of TFBS to form 14dTFBS_a revealed the importance of multiple TFBS for enhancer activity and a threshold requirement of more than ten TFBS. Error bars represent standard deviation, groups determined by one-way ANOVA to be significantly different (P <0.05) are labelled with different letters. B) Enhancers that regulate *Sox2* or *Med13l* contain multiple conserved TFBS (top) which are required for activity as demonstrated by TFBS mutagenesis (bottom). TFBS indicated in blue were required for enhancer activity. TFBS indicated in grey were not modified, yellow indicates TFBS found not to be required for activity. Significant differences compared to the wild type (WT) sequence are indicated by * P<0.05, or *** P<0.001, ns = not significant. Error bars represent standard deviation. C) Transcription factor bound regions in mouse ESC with >10 TFBS have significantly higher enrichment of H3K27ac and increased DNase I sensitivity (DNase-seq) compared to transcription factor bound regions with 8-10 and 1-7 TFBS in the 700bp sequence window. Groups determined by one-way ANOVA to be significantly different (P <0.0001) are labelled with different letters. Dashed line indicates the average for the 1-7 TFBS group.

TFBS removal from synthetic enhancers supported the hypothesis that TFBS diversity, above a threshold, confers enhancer activity. To investigate this threshold model in endogenous enhancers, and the TFBS repertoire we identified as enriched in CHEF sequences, we mutated identified conserved TFBS in two validated ESC enhancers [*Sox2* and *Med13l* enhancers, (Moorthy et al., 2017; Zhou et al., 2014)], both of which are bound by six or more of the core pluripotency regulatory transcription factors (Figure S4A/B). In both enhancers, we determined that at least nine of the conserved TFBS, enriched in CHEF regions, are required for enhancer activity (Figure 5B). Interestingly, although the *Sox2* enhancer contained eight different conserved TFBS confirmed to be important for activity, co-mutation of only two different TFBS (TFAP2C and SP1) completely abolished enhancer activity (Figure 5B). These data revealed that in addition to TFBS bound by the nine pluripotency transcription factors initially considered, additional TFBS (TFAP2C, SP1, ZNF263, GBX2, ZIC3, TEAD4, and FOXH1) are important for enhancer activity confirming that there is a large repertoire of TFBS that contribute to enhancer activity in pluripotent cells.

To determine if the TFBS diversity threshold model can identify regions of the genome with active enhancer features in mouse ESCs, we compared H3K27ac flanking the >18,000 transcription factor bound regions, to the number of different TFBS contained within each sequence, considering the 70 different TFBS enriched in CHEF regions (Table S2). This analysis revealed sequences with a greater number of different CHEF-enriched TFBS display significantly higher H3K27ac (P<0.0001, ANOVA, Figure 5C). Similarly, we found that regions containing an increased number of CHEF-enriched TFBS display significantly increased sensitivity to DNaseI (P<0.0001, ANOVA, Figure 5C). Although H3K27ac and sensitivity to DNaseI have been shown to correlate with enhancer activity these features can be observed at non-functional regions as shown by recent ChIP-STARR-seq and FAIRE-STARR-seq analysis (Barakat et al., 2018; Chaudhri et al., 2019). As the ability to drive transcription of a nearby gene is a better measure of enhancer activity, we examined the regulatory activity of these three TFBS classes (1-7, 8-10, and >10 CHEF-enriched TFBS) by comparing the expression of genes within a 200kb window. We determined that genes with a >10 TFBS region within 200kb displayed significantly increased gene expression compared to both the 1-7 and 8-10 CHEF-enriched TFBS groups (P<0.0001, ANOVA, Figure S5). Together these data indicate that highly heterotypic sequences containing a strong consensus for more than ten TFBS from the CHEF-enriched pluripotency repertoire are likely to be strong enhancers in ESCs.

## DISCUSION

Based on these data we propose a regulatory code for naïve pluripotent stem cells that requires more than ten TFBS from the repertoire of 70 TFBS enriched in CHEF sequences. Transcription factors that bind this expanded TFBS repertoire are likely to be important regulators in pluripotent cells which could drive more efficient reprogramming in combination with conventional reprogramming factors. In support of this, SP5, TFAP2C, GBX2, E2F3, ZIC3, FOXP1/FOXO1, ZFX and PRDM14 have recently been revealed as important regulators of pluripotency, self-renewal or reprogramming (Harel et al., 2012; Lim et al., 2007; Pastor et al., 2018; Seki, 2018; Tai and Ying, 2013; Tang et al., 2017, 2016; Zhang et al., 2011a).

TFBS sequence diversity above a threshold could facilitate recruitment of the large protein complexes required for transcriptional activation and elongation in mammalian cells. The mammalian mediator complex consists of 33 subunits and is required for RNA polymerase II transcription and gene expression in ESCs (El Khattabi et al., 2019). Furthermore, DNA sequences containing OCT4 TFBS act as a scaffold for phase-separated condensation of the mediator component MED1 in the nucleus (Shrinivas et al., 2019). In addition, within the set of transcription factors that bind the TFBS we identified as important for enhancer activity in pluripotent cells, individual transcription factors interact with different subcomponents of the mediator complex. SOX2 associates with MED22, NANOG with MED12, FOXJ3 with MED4 and 23, SMAD1 with MED6, 15, and 24, and ESRRB associates with several of the mediator complex proteins (Warde-Farley et al., 2010). Mediator recruitment could be facilitated by sequences that bind multiple transcription factors, each able to associate with different components of the complex.

Although examples exist where regulatory grammar is critical for activity as proposed by the enhanceosome model (Thanos and Maniatis, 1995), our findings indicate TFBS diversity above a threshold has a greater impact on enhancer activity. Previous studies using synthetic sequences support the observation that heterotypic sequences containing 4 different TFBS have increased regulatory activity compared to sequences with fewer different TFBS (Fiore and Cohen, 2016; King et al., 2020). In addition, two studies assaying native sequences containing 4-6 different TFBS found only 28-37% of these possessed enhancer activity suggesting additional TFBS present in the active sequences confer activity (King et al., 2020; Lloret-Fernández et al., 2018). The FOXP1 TFBS, a member of our pluripotency TFBS repertoire, was identified as one of the TFBS enriched in the genomic sequences displaying enhancer activity in ESCs by King et al. 2020. We have now shown that more than ten different TFBS are required for activity equivalent to a native enhancer, and that these TFBS can be drawn randomly from a large repertoire.

The observation that there is a large repertoire of TFBS that can be flexibly used to support enhancer activity without the need for optimized regulatory grammar explains the difficulty in identifying a specific regulatory code in mammalian genomes. A recent study in B-cells using STARR-seq coupled with FAIRE-seq identified 43 TFBS significantly enriched in B-cell enhancers (Chaudhri et al., 2019). Comparing these B-cell TFBS with the pluripotent cell TFBS repertoire we identified, revealed only 14% overlap, suggesting a large cell-type specific TFBS repertoire confers context dependent enhancer activity. Similarly, MPRA analysis revealed that enhancer activity of PPARy bound regions in adipocytes depends on additional flanking TFBS from a repertoire of 33 TFBS (Grossman et al., 2017), supporting our findings. We also determined that there is flexibility in TFBS usage from this larger tissue-specific repertoire and that this can bypass the need for a specific individual TFBS, even the OCT4:SOX2 co-motif which binds the OCT4, SOX2 and NANOG master regulators of pluripotency. This finding is supported by a recent study showing that OCT4 can be excluded from the set of reprogramming transcription factors without limiting reprogramming efficiency (Velychko et al., 2019). Our work also reinforces the idea that master-regulator transcription factors are an over-simplification and instead these regulators are part of a larger and complex ‘Gene Regulatory Network’ where inputs from all transcription factors in the network are similarly important (Singh et al., 2014). Deciphering the TFBS repertoire in pluripotent stem cells supported the design of short synthetic enhancers which allow for fine-tuned control of gene expression in biotechnology and regenerative medicine contexts. Determining the repertoire of specific TFBS important for enhancer activity in different cell types will inform a more mechanistic understanding of how specific enhancer SNPs affect gene expression and are linked to disease.

### Experimental Procedures

#### Analysis of conserved transcription factor bound regions

Transcription factor ChIP-seq data for mouse and human was obtained from ESCODE in the CODEX database and GEO (Table S5)(Sánchez-Castillo et al., 2014; Tsankov et al., 2015). Mouse transcription factor bound regions were identified from 16 files representing binding of nine transcription factors (OCT4, SOX2, NANOG, KLF4, KLF2, ESRRB, SMAD1, STAT3, and TCFCP2L1). Human transcription factor bound regions were obtained from 9 files representing binding of seven transcription factors (OCT4, SOX2, NANOG, KLF5, SMAD1, STAT3 and TCF4). Overlapping transcription factor bound regions were identified within 1kb windows. Mouse-human syntenic regions were identified using liftOver (−minMatch=0.1)(Kuhn et al., 2013). The human sequences were extended by 1kb from the mid-point to identify conserved transcription factor binding using BEDTools (Quinlan and Hall, 2010). ChIP-seq data was retrieved for H3K27ac (GSE47949)(Wamstad et al., 2012) and EP300 (GSE24164)(Creyghton et al., 2010) in mouse ESC or H3K27ac in naïve human ESCs (GSE52824)(Gafni et al., 2013). The reads were mapped to respective genomes (mm10 and hg19) using Bowtie(Langmead et al., 2009). RPKM values for H3K27ac ChIP/input were calculated for 2kb regions surrounding transcription factor bound region midpoints. Venn diagrams were created using ‘eulerr’ in R (Micallef and Rodgers, 2014).

To focus on conserved distal regulatory regions the 23,830 transcription factor bound regions in mouse that aligned to the human genome after excluding promoters (transcription start site, TSS −2 kb) were clustered into 9 clusters. The H3K27ac ChIP-Seq signal was assumed to follow a Gaussian distribution, whereas binding of each transcription factors were assumed to follow Bernoulli distributions. The sums of mouse & human transcription factors (TF sum in Figure 1) were assumed to follow Poisson distributions. To optimize parameters of the mixture model, we used the expectation maximization algorithm, which requires initial parameter values. We used Behrouz Babaki’s implementation of constrained k-means (https://github.com/Behrouz-Babaki/COP-Kmeans) (Wagstaff et al., 2001) to generate initial clusters. Using the ‘pomegranate’ package, ‘IndependentComponentsDistribution’ was used to create a multivariate distribution using the initial clusters’ parameters and finally the ‘GeneralMixtureModel’ function was used to fit and predict the clusters of enhancers (Schreiber, 2018). For the cluster visualization a heatmap was generated using the ‘seaborn’ package (Qalieh et al., 2017), in which each feature is scaled from 0 to 1. The cluster output with the lowest Bayesian information criterion (BIC) was chosen as the final cluster.

#### TFBS conservation and enrichment analysis

TFBS analysis was carried out for sequences within each cluster and random intergenic regions selected from regions not bound to a transcription factor in mouse or human ESCs. Any region overlapping an exon or 2kb upstream of a TSS was removed as these sequences are often highly conserved. Using 700bp of sequence centered on the midpoint of the transcription factor bound region (or sequence midpoint for non-transcription factor bound regions) in the mouse genome, the corresponding syntenic regions in the human, rhesus macaque, mouse, rat, cow and pig genomes were identified using liftOver. After liftOver the sequence was expanded to 2kb prior to multi-sequence alignment (MSA). MSA was performed using Mafft (Katoh and Standley, 2013) E-INS-I (Iterative refinement method) with the ‘adjustdirectionaccurately’ feature. MSA was trimmed on both 5’ and 3’ ends to best match the 700bp mouse reference sequence with respective aligned sequences from the other five species. The mouse sequence was scanned for TFBS with a relative profile threshold of 90% for all 519 non-redundant JASPAR profiles by using ‘searchSeq’ in ‘TFBSTools’ (Tan and Lenhard, 2016) and the ‘JASPAR2016’ (Mathelier et al., 2016) package in R. For purifying selection pressure analysis of the CHEF cluster, putative sites with a score >11 for each JASPAR PWM were evaluated. Human-mouse percent identity for each TFBS was calculated using TFBSTools identified motifs in the mouse sequence. The corresponding sequences for human were then identified in the MSA. TFBS conservation was calculated as average percentage identity between mouse and human of all motifs for each individual PWM. Scrambled PWM for all 519 JASPAR profiles were created using ‘permuteMatrix’ in ‘TFBSTools’ (Tan and Lenhard, 2016) and then those scrambled PWM were scanned in the mouse sequence for CHEF regions and mouse-human percentage identity was calculated as described above to identify scrambled TFBS conservation.

The 700 bp trimmed six species MSA for each of the regions in the 9 clusters and random regions was used as input for MotEvo (Arnold et al., 2012) with mouse as the reference sequence. Input parameters for MotEvo included nucleotide percentage and averaged phylogenetic tree for all six species which were calculated using the MSA of CHEF regions. Phylogenetic trees were generated using PhyML (Guindon et al., 2010) for those MSA with liftOver sequences in all six species. MotEvo analysis was performed using refspecies Mouse; Mode: TFBS; EMprior: 0; markovorderBG: 0; minposterior: 0.2. Motevo output resulted in posterior probabilities for each of the 349 redundant PWM belonging to expressed transcription factors in ESCs (log RPKM≥0.025, RNA-Seq using GEO accession: GSE63831), which were used as features in machine learning to distinguish CHEF sequences from the NANOG cluster. To rank and select important features, LASSO (Least Absolute Shrinkage and Selection Operator) was used with the Scikit-learn (Pedregosa et al., 2011) Python module. LassoLarsCV with 10-fold cross validation was used to identify the parameter (−log[alpha]) which generated the highest classification accuracy which was then used for feature selection.

To determine the number of conserved different TFBS present in cluster regions the number of different MotEvo identified motifs, from the set of 349 TFBS, were counted in each sequence. To determine if conserved TFBS are present in addition to any corresponding to the 9 transcription factors used for cluster analysis, the number of different MotEvo identified motifs, from a reduced set of 309 TFBS, were counted in each sequence.

349 TFBS belonging to transcription factors expressed in ESC were reduced to 137 TFBS significantly different from each other (Pearson correlation coefficient <0.65) which included 70 TFBS enriched in CHEF (LASSO>0) using TFBSTools’ (Tan and Lenhard, 2016). This set of enriched TFBS was used to identify the number of different TFBS present within a 700bp window for all transcription factor bound regions (>18,000). Motifs with score >11 as described above were identified using TFBSTools. The relationship between the increasing number of different TFBS from the enriched TFBS set and H3K27ac/DHS (GSE37074) in the 2kb flanking regions was analysed.

#### Site-directed mutagenesis

For the analysis of conserved TFBS in the *Sox2* and *Med13l* enhancers, the two nucleotides in the motif with the highest score in the consensus were mutated to non-consensus nucleotides. The effect of the mutation was tested using TFBSTools to confirm that it disrupted the reference TFBS without creating another TFBS for an ESC expressed TF. Mutations in enhancer sequences in pGL4.23 vector were introduced using the QuickChange Lightening kit (Agilent) with the primers described in Table S6. For the gain of function analysis in the *Sall1* MTL52 sequence we focused on the smaller core region bound by six transcription factors (MTL52core). We identified motifs with the closest match to the desired TFBS, as these all had a low consensus match score, the identified motifs were mutated in the MTL52core sequence using Gibson Assembly to produce the consensus sequence for ESRRB, TFCP2L1, and SMAD3 (+E+T+S). In the +E+T+S sequence, ESRRB motif was mutated to non-consensus nucleotides preventing binding of any other transcription factor at this region to create +T+S. Another gain of function construct +T+S+E2F1 was created by adding a strong consensus motif for E2F1 to the +T+S construct using the QuickChange Lightening kit (Agilent). For the gain of function TFBS analysis, a maximum of six mutations were introduced, using the primers described in Table S6, to convert a non-consensus motif into a strong consensus motif for the desired TFBS.

#### Cell culture and CRISPR deletion

F1 mouse ES cells (*M. musculus*^129^ × *M. castaneus*, obtained from Barbara Panning) were cultured on 0.1% gelatin-coated plates in ESC media (DMEM containing 15% FBS, 0.1 mM MEM nonessential amino-acids, 1 mM sodium pyruvate, 2 mM GlutaMAX, 0.1 mM 2-mercaptoethanol, 1000 U/mL LIF, 3 µM CHIR99021 [GSK3β inhibitor; Biovision], and 1 µM PD0325901 [MEK inhibitor; Sigma Aldrich]), to maintain the naïve state. Cas9 targeting guides flanking *Sall1* proximal transcription factor bound regions were selected (Table S7). Only gRNAs predicted to have no off-target binding in the mouse genome were chosen. Guide RNA plasmids were assembled in the gRNA empty vector (Addgene, ID#41824) using the protocol described by Mali et. al. (2013)(Mali et al., 2013) and confirmed by sequencing. Following the protocol described in Moorthy et. al. (2016)(Moorthy and Mitchell, 2016), 5 µg each of the indicated gRNAs targeting the 5’ and 3’ ends of the transcription factor bound region were transfected in ESCs with pCas9_GFP (Addgene, ID#44719)(Ding et al., 2013) using the Neon Transfection System (Life Technologies). FACS sorting 48 hr post-transfection isolated GFP-positive cells, which were collected and seeded on 10 cm gelatinized culture plates. Genotyping of enhancer deletions was done by qPCR with allele-specific primers. All deletions were confirmed by sequence analysis using primers 5′ of and 3′ from the gRNA target sites; SNPs within the amplified product confirmed the genotype of the deleted allele. Gene expression was monitored by RT-qPCR using allele-specific primers (Sall1_129, Sall1_Cast) that distinguish 129 from *castaneus* alleles (Table S6). The standard curve method was used to calculate expression levels, with mouse F1 genomic DNA used to generate the standard curves.

#### Synthetic sequences, enhancers and luciferase assay

Synthetic enhancer sequences as described in Table S4 were generated as GenPart DNA Fragments (GenScript). These sequences contained 2bp ‘CC’ spacers between each TFBS. All the 349 TFBS belonging to ES expressed transcription factors were clustered to 137 diverse TFBS sets with one or more TFBS in each set with significantly similar PWM to each other (Pearson correlation coefficient ≥0.65). Based on the LASSO coefficients of the representative TFBS from each of these 137 diverse TFBS sets, they were classified into three classes (LASSO>0, LASSO=0, LASSO<0). For the creation of 14dTFBSa-c, all 14 unique TFBS were chosen from the LASSO>0 class, while for 14dTFBSd-g an increasing number of TFBS from the LASSO<0 class were chosen. Using the conserved motifs for selected TFBS identified from the MotEvo analysis, the distance and orientation between each pair of TFBS was averaged over all CHEF sequences to identify the average co-occurrence distance and preferred orientation between each TFBS pair. If more than one motif per TFBS were present, the orientation present in ≥66% motifs was chosen as reference otherwise it was excluded. Significance of orientation bias was calculated by Binomial Test, with the null hypothesis being no bias, 50% occurrence probability for either orientation.

To evaluate enhancer activity, synthetic sequences or native transcription factor bound regions were cloned downstream of the firefly luciferase gene at the *NotI* site in the pGL4.23 vector containing an *Oct4* minimal promoter as described in Moorthy et. al. (Moorthy et al., 2017). Mouse F1 ESCs were seeded on gelatin coated 96-well plates at a density of 10,000 cells per well. After 24 hours the cells were co-transfected using jetPRIME (VWR International) with pGL4.23 vectors and a *Renilla* luciferase encoding plasmid (pGL4.75) at a 50:1 molar ratio. Renilla plasmid is transfected as control for transfection efficiency in each well. ESC media was replaced with fresh ESC media after 24 hours. Luciferase activity was assayed 48 hours post transfection using a dual luciferase reporter assay (Promega) and the firefly/Renilla ratio was measured on the Fluoroskan Ascent FL plate reader. All experiments were done in ≥3 biological replicates with each experiment having ≥3 technical transfection replicates.

## Author Contributions

G.S. and J.A.M. conceived the project, designed the experiments, and analyzed all experimental data. A.M.M. contributed to the design and analysis for computational approaches. G.S. performed the experiments in Figures 1A-E, 2A-B, 3A-C, 5A-C, S1A-D, S2A-B, S4A-B and S5. S.M. performed the experiments in Figures 3A and 4D.

S.D.M. performed the experiments in Figures 4B and S3B. R.Z. performed the experiments in Figure 5B. T.M. performed the experiments in Figure 1B. R.T. performed the experiments in Figure 4D. G.S. and J.A.M. wrote the manuscript, which was approved by all co-authors.

## ACKNOWLEDGMENTS

We would like to thank all the members of the Mitchell and Moses labs for helpful discussions. Thank you to Virlana M Shchuka who performed the experiment in Figure 4C. This work was supported by Natural Science and Engineering Research Council of Canada, the Canada Foundation for Innovation, and the Ontario Ministry of Research and Innovation (operating and infrastructure grants held by J.A.M.). Studentship funding to support this work was provided by Connaught International Scholarships and Ontario Graduate Scholarship (G.S.). Salary support for S. D. M. was provided by Canadian Institutes of Health Research Project Grant (FRN 153186).

