## Supplemental Data for "A flexible repertoire of transcription factor binding sites and diversity threshold determines enhancer activity in embryonic stem cells"

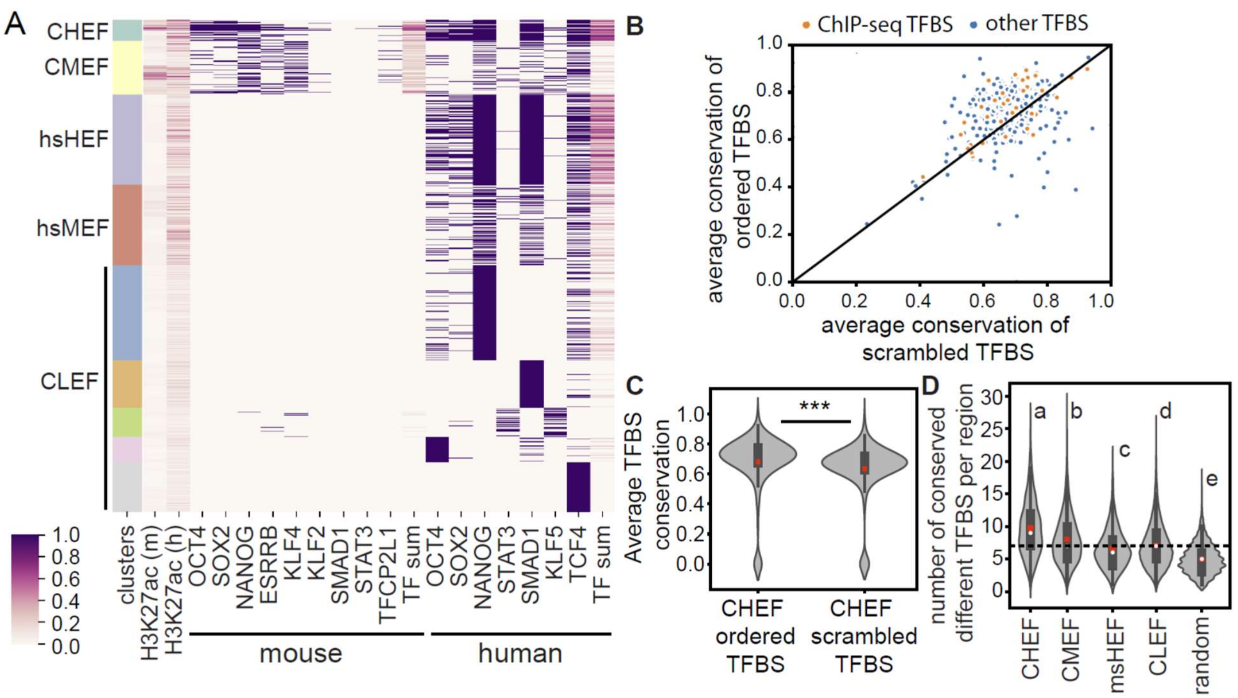


**Supplemental Figure S1: Mouse-human conserved high enhancer feature regions (CHEF) contain conserved TFBS.** A) Clustering of TF bound regions in human ESCs, using H3K27ac histone modification in human, TF binding and H3K27ac histone modification at associated mouse regions. Clusters are labeled as CHEF, conserved medium enhancer feature (CMEF), human specific high enhancer feature (hsHEF), human specific medium enhancer feature (hsMEF), and conserved low enhancer feature (CLEF) regions. B) Average TFBS conservation for ordered TFBS compared to scrambled TFBS in mouse CHEF regions. Each dot represents a different specific TFBS. The majority of the ordered TFBS are above the diagonal line indicating they are more conserved than the scrambled TFBS. C) Average TFBS conservation in mouse CHEF regions is significantly higher for ordered TFBS compared to scrambled TFBS. D) CHEF cluster regions in mouse contain an increased number of conserved different TFBS compared to other clusters and random non-TF bound regions after removal of all TFBS bound by ChIP-seq TFs. Groups determined by one-way ANOVA to be significantly different ($P<0.05$) are labelled with different letters.

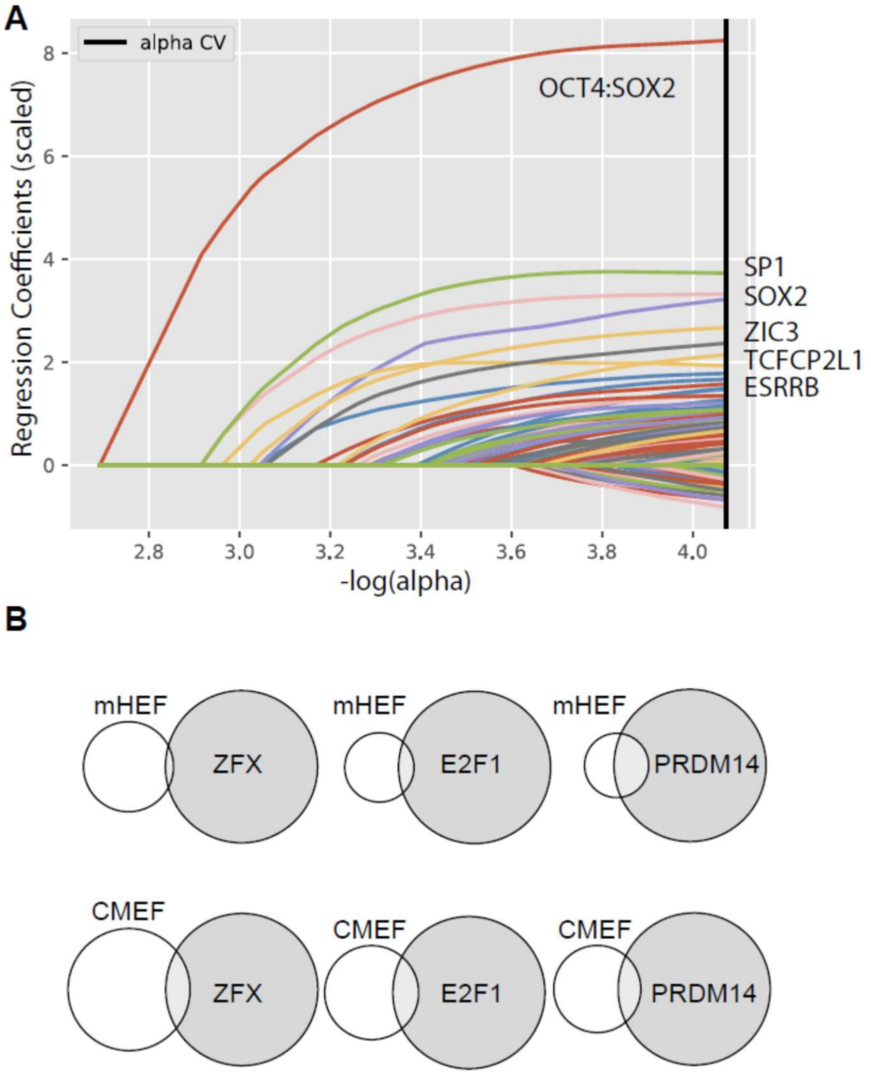


**Supplemental Figure S2: TFBS enrichment in CHEF regions and those mutated by site directed mutagenesis (SDM).** A) Regression coefficient progression for LASSO paths. B) Overlap of mouse CMEF and msHEF regions with the EP300 bound regions and ChIP-seq for PRDM14, E2F1, or ZFX.

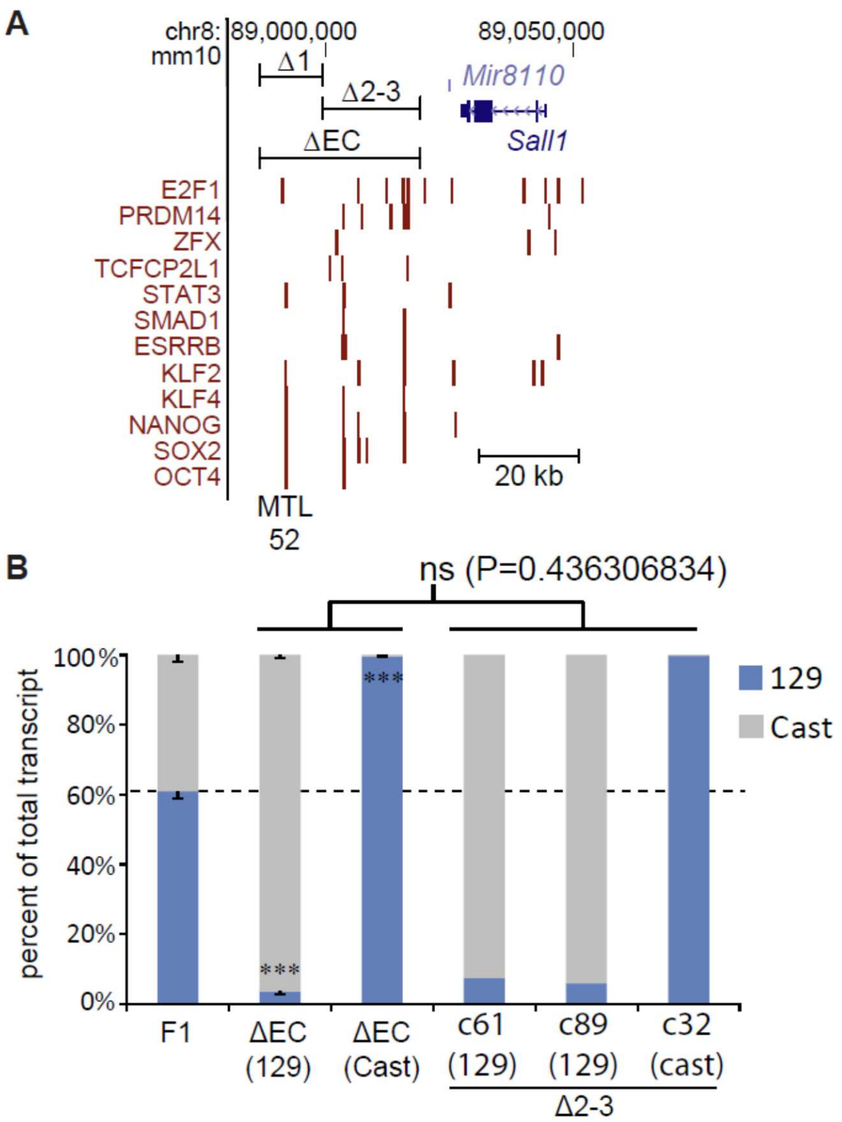


**Supplemental Figure S3: The multiple transcription factor bound region 52 kb downstream of *Sall1* (MTL52) does not drive transcription of *Sall1* in ESCs.** A) Schematic representation of the *Sall1* locus in the mouse genome. Transcription factor bound regions from ChIP-seq (red bars) are displayed on the mm10 assembly of the University of California at Santa Cruz (UCSC) Genome Browser. CRISPR deleted regions (ΔEC, Δ1, Δ2-3) are displayed. The location of MTL52 in the Δ1 region is indicated. B) *Sall1* expression in wild type F1 clones (F1) compared to clones with the indicated deletion. Allele-specific primers detect 129 or Cast RNA in RT-qPCR. Expression for each allele is shown relative to the total. Error bars represent SEM. Significant differences from the F1 values are indicated by *** $P < 0.001$ for the EC deletion replicates. Deletion of the entire EC dramatically reduces expression of the linked *Sall1* allele. Deletion of the Δ2-3 region similarly reduces expression of the linked *Sall1* allele in 3 separate clones (c61, c89, c32). Comparison of the percent reduced transcription for ΔEC (n=8) compared to Δ2-3 (n=3) revealed no significant difference (ns), indicating MTL52 does not significantly contribute to *Sall1* transcription in ESs.

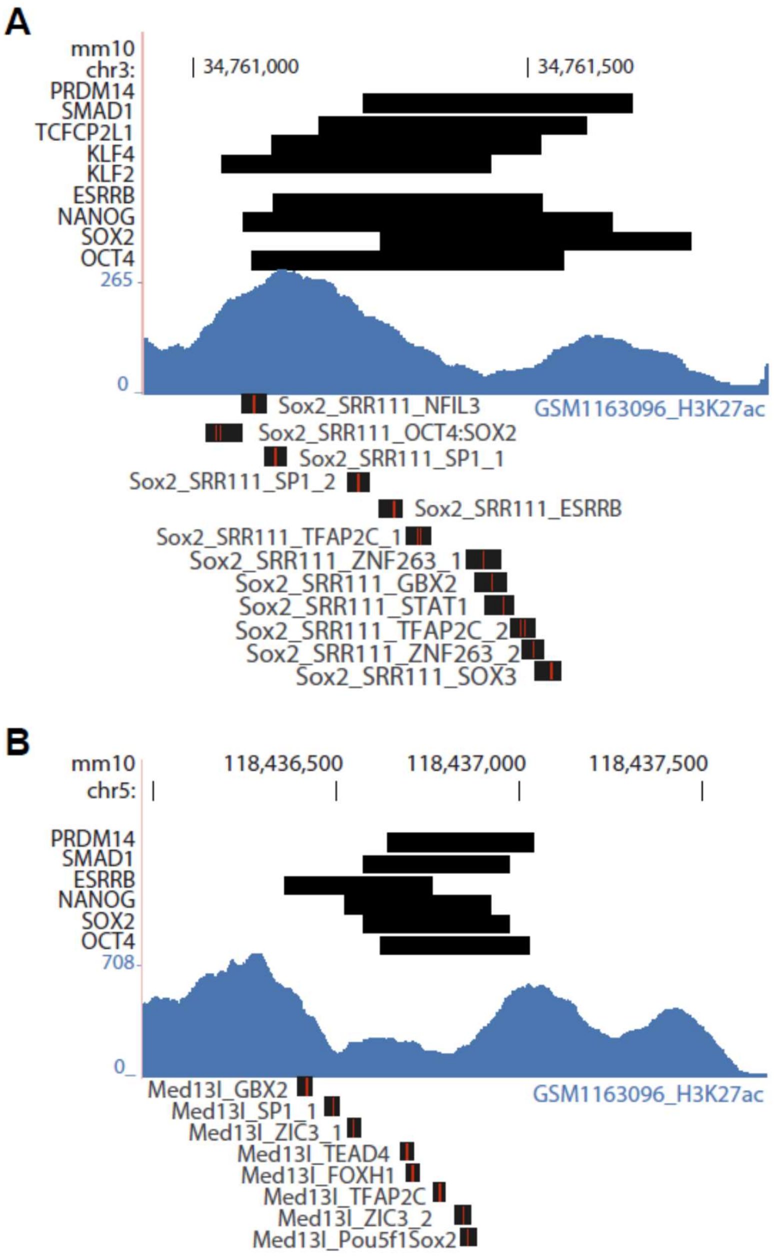


**Supplemental Figure S4: Mutations in the Sox2 or Med13l enhancers.** Schematic representation of the *Sox2* (A) or *Med13l* (B) enhancer in the mouse genome. Transcription factor bound regions from ChIP-seq (black bars at top) are displayed on the mm10 assembly of the University of California at Santa Cruz (UCSC) Genome Browser. Mutations introduced by SDM are indicated by red lines in black bars below. H3K27ac ChIP-seq data from ESCs is displayed in blue.

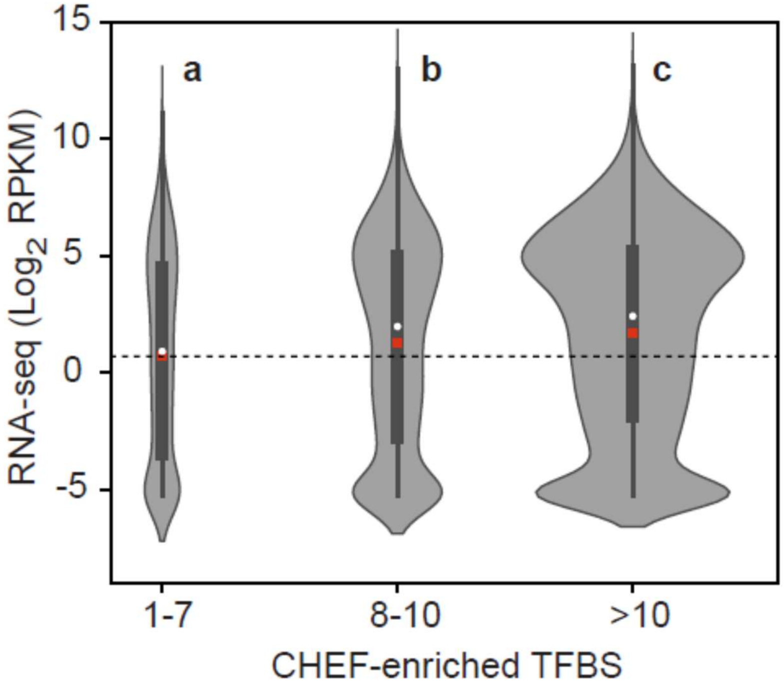


**Supplemental Figure S5: Genes within 200kb of TF bound regions containing an increased number of CHEF-enriched TFBS are expressed at higher levels.** Genes were separated into 3 groups; those with a >10 TFBS region within 200kb, an 8-10 TFBS or 1-7 TFBS region within 200kb. Groups determined by one-way ANOVA to be significantly different (P <0.0001) are labelled with different letters. Dashed line indicates the average for the 1-7 TFBS group.

**Table S1: Clustering of transcription factor bound regions in mouse and human ESCs.**
Data provided in a separate file.

**Table S2: LASSO coefficients for TFBS enriched or depleted in CHEF compared to the NANOG bound cluster.**
Data provided in a separate file.

**Table S3: Orientation preferences in CHEF sequences.**
Data provided in a separate file.

**Table S4: Synthetic enhancer sequences.**
Vector sequence added for assembly is underlined.

<table>
<tr><td>14OS</td><td>OCT4:SOX2, OCT4:SOX2, OCT4:SOX2, OCT4:SOX2, OCT4:SOX2, OCT4:SOX2, OCT4:SOX2, OCT4:SOX2, OCT4:SOX2, OCT4:SOX2, OCT4:SOX2, OCT4:SOX2, OCT4:SOX2, OCT4:SOX2</td></tr>
<tr><td colspan="2"><u>GGTCTGACAGCGGCCGCACTTGTGCCTGAACACCATATC</u>CCctttgttatgcaaatCCctttgttatgcaaatCCctttgttatgcaaatCCctttgttatgcaaatCCctttgttatgcaaatCCctttgttatgcaaatCCctttgttatgcaaatCCctttgttatgcaaatCCctttgttatgcaaatCCctttgttatgcaaatCCctttgttatgcaaatCCctttgttatgcaaatCCctttgttatgcaaatCCctttgttatgcaaatCC<u>GTTTTCTTGGCAATCGACTCTCGCGGCCGCAAATGCTAA</u></td></tr>
<tr><td>ksOE(4TFBS) (longer)</td><td>klf4, sox2, Oct4, Esrrb</td></tr>
<tr><td colspan="2"><u>GGTCTGACAGCGGCCGCACTTGTGCCTGAACACCATATC</u>AGCTACCGGCCCCGCCCCGTAGCTACGAAACAATGAGCGTAGCTACGGGATGCTAATCGTAGCTACGTTCAAGGTCACGT<u>GTTTTCTTGGCAATCGACTCTCGCGGCCGCAAATGCTAA</u></td></tr>
<tr><td>sOKE(4TFBS) (longer)</td><td>sox2, Oct4, Klf4, Esrrb</td></tr>
<tr><td colspan="2"><u>GGTCTGACAGCGGCCGCACTTGTGCCTGAACACCATATC</u>AGCTACGAAACAATGAGCGTAGCTACGGGATGCTAATCGTAGCTACGGGGCGGGGCCGGTAGCTACGTTCAAGGTCACGT<u>GTTTTCTTGGCAATCGACTCTCGCGGCCGCAAATGCTAA</u></td></tr>
<tr><td>ksOE(4TFBS) (CC)</td><td>klf4, sox2, Oct4, Esrrb</td></tr>
<tr><td colspan="2"><u>GGTCTGACAGCGGCCGCACTTGTGCCTGAACACCATATC</u>CCCGGCCCCGCCCCCCGAAACAATGAGCCCGGGATGCTAATCCCGTTCAAGGTCACCC<u>GTTTTCTTGGCAATCGACTCTCGCGGCCGCAAATGCTAA</u></td></tr>
<tr><td>sOKE(4TFBS) (CC)</td><td>sox2, Oct4, Klf4, Esrrb</td></tr>
<tr><td colspan="2"><u>GGTCTGACAGCGGCCGCACTTGTGCCTGAACACCATATC</u>CCGAAACAATGAGCCCGGGATGCTAATCCCGGGGCGGGGCCGCCGTTCAAGGTCACCC<u>GTTTTCTTGGCAATCGACTCTCGCGGCCGCAAATGCTAA</u></td></tr>
<tr><td>14dTFBS_a</td><td>Zic3, OCT4:SOX2, Tfcp2, Smad3, Sp1, ZNF263, KLf4, Ahr::Arnt, Foxh1, Foxj3, Esrrb, STAT1, TFAP2C, TEAD4</td></tr>
<tr><td colspan="2"><u>GGTCTGACAGCGGCCGCACTTGTGCCTGAACACCATATC</u>CCggcccccgctgtgcCCctttgttatgcaaatCCAAACCGGTTTCCgtctagacaCCccccgcccccCCggaggaggaggaggaggaggaCCtaaaggaaggCCgcagacaagcagcggCCtccaatccacaCCataaagtaaacaaacacCCagctcaaggtcaCCagaaaatgaaactgCCtgccctagggcaCCcacattccatCC<u>GTTTTCTTGGCAATCGACTCTCGCGGCCGCAAATGCTAA</u></td></tr>
<tr><td>14dTFBS_b</td><td>FOXP1, STAT1:STAT2, ZFP263, SMAD3, ZIC3, LIN54, ESRRB, TFCP2, PRDM1, MLXIP, E2F3, RREB1, TEAD3, TCF7L2</td></tr>
<tr><td colspan="2"><u>GGTCTGACAGCGGCCGCACTTGTGCCTGAACACCATATC</u>CCataaagtaaacaaacacCCgtagtttcatttcccCCggaggaggaggaggaggaggaCCgtctagacaCCggcccccgctgtgcCCatttgaattCCagctcaaggtcaCCAAACCGGTTTCCagaaag</td></tr>
</table>

<table>
<tr><td colspan="2">tgaaagttaCCgcacgtgtCCctcccgcccccgcccCCacccaaaccacccccacacaCCcacattccatCCgaagttcaaaggaaCCGTTTTCTTGGCAATCGACTCTCGCGGCCGCAAATGCTAA</td></tr>
<tr><td>14dTFBS_c</td><td>TFCP2, TFAP2C, SMAD3, ESRRB, LIN54, ZIC3, ZFX, ZBTB7A, FOXH1, PRDM1, MLXIP, TCF7L2, FOXP1, ZFP263</td></tr>
<tr><td colspan="2">GGTCTGACAGCGGCCGCACTTGTGCCTGAACACCATATCCCAAACCGGTTTCCtgccctagggcaCCgtctagacaCCagctcaaggtcaCCatttgaattCCggcccccgctgtgcCCcccgccgcggcctgCCggcgaccacagaCCtccaatccacaCCagaaagtgaaagttaCCgcacgtgtCCgaagttcaaaggaaCCataaagtaaacaaacacCCggaggaggaggaggaggaggaCCGTTTTCTTGGCAATCGACTCTCGCGGCCGCAAATGCTAA</td></tr>
<tr><td>14dTFBS_d</td><td>HIC2, FOXD1, Myc, MAFK, Hic1, TFEB, TP53, ZNF410, HINFP, HLF, TFAP2C(var.2), Bhlhe40, JUNB, Stat4</td></tr>
<tr><td colspan="2">GGTCTGACAGCGGCCGCACTTGTGCCTGAACACCATATCCCatgcccaccCCgtaaacatCCccatgtgcttCCaagactcagcaatttCCatgccaaccCCatcacgtgacCCctggacatgcctgtgcctgtCCtgcatcccataataactCCctacgtccgcCCggttacataattCCagcctcaggcaCCctcacgtgcacCCggatgactcatCCtttccaggaaataaCCGTTTTCTTGGCAATCGACTCTCGCGGCCGCAAATGCTAA</td></tr>
<tr><td>14dTFBS_e</td><td>JUNB, FOSL1, Hic1, Stat4, Ddit3::Cebpa, TFAP2C(var.2), HLF, TFEB, TP53, GLI2, Bhlhe40, PBX1, Klf12, MAX</td></tr>
<tr><td colspan="2">GGTCTGACAGCGGCCGCACTTGTGCCTGAACACCATATCCCggatgactcatCCggtgactcatgCCatgccaaccCCtttccaggaaataaCCggatgcaatcccCCagcctcaggcaCCggttacataattCCatcacgtgacCCacatgccctggcatgCCgcgaccacactgCCctcacgtgcacCCccatcaatcaaaCCgaccacgccctttctCCaagcacatggCCGTTTTCTTGGCAATCGACTCTCGCGGCCGCAAATGCTAA</td></tr>
<tr><td>14dTFBS_f</td><td>SREBF2, MEF2D, Bhlhe40, Stat4, JUNB, TFEB, ETV5, REST, HIC2, SMAD2::SMAD3::SMAD4a, MTF1, RARA, MAFK, Klf12</td></tr>
<tr><td colspan="2">GGTCTGACAGCGGCCGCACTTGTGCCTGAACACCATATCCCatggggtgatCCactataaatagcCCctcacgtgcacCCtttccaggaaataaCCggatgactcatCCatcacgtgacCCaccggaagtgCCgcgctgtccctggttctgaCCatgcccaccCCtgtctgtcacctCCtttgcacacggcacCCaaggtcagaagttcaaggCCaagactcagcaatttCCgaccacgccctttctCCGTTTTCTTGGCAATCGACTCTCGCGGCCGCAAATGCTAA</td></tr>
<tr><td>14dTFBS_g</td><td>Ddit3::Cebpa, EN1, GLI2, MAX, TCF4, SMAD2::SMAD3::SMAD4a, SREBF2, PBX1, Hic1, Gmeb1, HIC2, FOXD1, Myc, MTF1</td></tr>
<tr><td colspan="2">GGTCTGACAGCGGCCGCACTTGTGCCTGAACACCATATCCCggatgcaatcccCCctaattagCCgcgaccacactgCCaagcacatggCCgcacctgctCCctgtctgtcacctCCatggggtgatCCccatcaatcaaaCCatgccaaccCCcagtttacgtaagaagaCCatgcccaccCCgtaaacatCCccatgtgcttCCtttgcacacggcacCCGTTTTCTTGGCAATCGACTCTCGCGGCCGCAAATGCTAA</td></tr>
<tr><td>14dTFBS</td><td>Zic3, OCT4:SOX2, Tfcp2, Smad3, Sp1, ZNF263, KLf4, Ahr::Arnt, Foxh1, Foxj3, Esrrb, STAT1, TFAP2C, TEAD4</td></tr>
<tr><td colspan="2">GGTCTGACAGCGGCCGCACTTGTGCCTGAACACCATATCCCggcccccgctgtgcCCctttgttatgcaaatCCAAACCGGTTTCCcgtctagacaCCccccgcccccCCggaggaggaggaggaggaggaCCtaaaggaaggCCgcagacaagcagcggCCtccaatccacaCCataaagtaaacaaacacCCagctcaaggtcaCCagaaaatgaaactgCCtgccctagggcaCCcacattccatCCGCGGCCGCAAATGCTAA</td></tr>
<tr><td>12dTFBS</td><td>Zic3, OCT4:SOX2, Tfcp2, Smad3, Sp1, ZNF263, KLf4, Ahr::Arnt, Foxh1, Foxj3, Esrrb, STAT1</td></tr>
<tr><td colspan="2">GGTCTGACAGCGGCCGCACTTGTGCCTGAACACCATATCCCggcccccgctgtgcCCctttgttatgcaaatCCAAACCGGTTTCCcgtctagacaCCccccgcccccCCggaggaggaggaggaggaggaCCtaaaggaaggCCgcagacaagcagcggCCtccaatccacaCCataaagtaaacaaacacCCagctcaaggtcaCCagaaaatgaaactgCCGCGGCCGCAAATGCTAA</td></tr>
<tr><td>10dTFBS</td><td>Zic3, OCT4:SOX2, Tfcp2, Smad3, Sp1, ZNF263, KLf4, Ahr::Arnt, Foxh1, Foxj3</td></tr>
</table>

| | |
|---|---|
| GGTCTGACAGCGGCCGCACTTGTGCCTGAACACCATATCCCggcccccgctgtgcCCctttgttatgcaaatCCAAACCGGTTTCCcgtctagacaCCccccgcccccCCggaggaggaggaggaggaggaCCtaaaggaaggCCcgcagacaagcagcggCCtccaatccacaCCataaagtaaacaaacacCCGCGGCCGCAAATGCTAA | |
| **7dTFBS** | Zic3, OCT4:SOX2, Tfcp2, Smad3, Sp1, ZNF263, KLf4 |
| GGTCTGACAGCGGCCGCACTTGTGCCTGAACACCATATCCCggcccccgctgtgcCCctttgttatgcaaatCCAAACCGGTTTCCcgtctagacaCCccccgcccccCCggaggaggaggaggaggaggaCCtaaaggaaggCCGCGGCCGCAAATGCTAA | |
| **5dTFBS** | Zic3, OCT4:SOX2, Tfcp2, Smad3, Sp1 |
| GGTCTGACAGCGGCCGCACTTGTGCCTGAACACCATATCCCggcccccgctgtgcCCctttgttatgcaaatCCAAACCGGTTTCCcgtctagacaCCccccgcccccCCGCGGCCGCAAATGCTAA | |
| **13dTFBS_rOri** | Zic3, OCT4:SOX2, Tfcp2, Smad3, Sp1, ZNF263, KLF4, Foxh1, Foxj3, Esrrb, STAT1, TFAP2C, TEAD4 |
| GGTCTGACAGCGGCCGCACTTGTGCCTGAACACCATATCCCggcccccgctgtgcCCctttgttatgcaaatCCAAACCGGTTTCCcgtctagacaCCccccgcccccCCggaggaggaggaggaggaggaCCtaaaggaaggCCtccaatccacaCCataaagtaaacaaacacCCagctcaaggtcaCCagaaaatgaaactgCCtgccctagggcaCCcacattccatCCGTTTTCTTGGCAATCGACTCTCGCGGCCGCAAATGCTAA | |
| **13dTFBS_rOri+NFYA (14TFBS)** | Zic3, OCT4:SOX2, Tfcp2, Smad3, Sp1, ZNF263, KLF4, NFYA, Foxh1, Foxj3, Esrrb, STAT1, TFAP2C, TEAD4 |
| GGTCTGACAGCGGCCGCACTTGTGCCTGAACACCATATCCCggcccccgctgtgcCCctttgttatgcaaatCCAAACCGGTTTCCcgtctagacaCCccccgcccccCCggaggaggaggaggaggaggaCCtaaaggaaggCCcgcagccaatcagcggCCtccaatccacaCCataaagtaaacaaacacCCagctcaaggtcaCCagaaaatgaaactgCCtgccctagggcaCCcacattccatCCGTTTTCTTGGCAATCGACTCTCGCGGCCGCAAATGCTAA | |
| **13dTFBS_pOri** | Zic3, OCT4:SOX2, Tfcp2, Smad3, Sp1, *znf263, klf4*, Foxh1, Foxj3, Esrrb, STAT1, TFAP2C, TEAD4 |
| GGTCTGACAGCGGCCGCACTTGTGCCTGAACACCATATCCCggcccccgctgtgcCCctttgttatgcaaatCCAAACCGGTTTCCcgtctagacaCCccccgcccccCCtcctcctcctcctcctccCCccttcctttaCCtccaatccacaCCataaagtaaacaaacacCCagctcaaggtcaCCagaaaatgaaactgCCtgccctagggcaCCcacattccatCCGTTTTCTTGGCAATCGACTCTCGCGGCCGCAAATGCTAA | |

**Table S5: Genome wide TF data used**

| GEO Accession | ChIP | Species | Purpose |
|---|---|---|---|
| GSM288355 | Esrrb | Mouse | Clustering |
| ERR440999 | Klf2 | Mouse | Clustering |
| ERR440998 | Klf2 | Mouse | Clustering |
| GSM288354 | Klf4 | Mouse | Clustering |
| GSM288345 | Nanog | Mouse | Clustering |
| GSM307140 | Nanog | Mouse | Clustering |
| GSM1090230 | Nanog | Mouse | Clustering |
| GSM288346 | Oct4_TF | Mouse | Clustering |
| GSM566277 | Oct4_TF | Mouse | Clustering |
| GSM307137 | Oct4_TF | Mouse | Clustering |
| GSM288348 | Smad1 | Mouse | Clustering |
| GSM288347 | Sox2 | Mouse | Clustering |
| GSM307138 | Sox2 | Mouse | Clustering |

| | | | |
|---|---|---|---|
| GSM1050291 | Sox2 | Mouse | Clustering |
| GSM288353 | Stat3 | Mouse | Clustering |
| GSM288350 | Tcfcp2l1 | Mouse | Clustering |
| GSM1505690 | Klf5 | Human | Clustering |
| GSM1505745 | Smad1 | Human | Clustering |
| GSM1505781 | Stat3 | Human | Clustering |
| GSM1505791 | Tcf4 | Human | Clustering |
| GSM1124071 | Nanog | Human | Clustering |
| GSM1124070 | Nanog | Human | Clustering |
| GSM1124069 | Sox2 | Human | Clustering |
| GSM1124068 | Sox2 | Human | Clustering |
| GSM1124067 | Oct4_TF | Human | Clustering |
| GSE52824 | H3K27ac | Human | Clustering |
| GSE47949 | H3K27ac | Mouse | Clustering |
| GSE24164 | EP300 | Mouse | MTL Validation |
| GSM288349 | E2F1 | Mouse | Enrichment Validation |
| GSM288352 | Zfx | Mouse | Enrichment Validation |
| GSM415050 | Mtf2 | Mouse | Enrichment Validation |
| GSM623989 | Prdm14 | Mouse | Enrichment Validation |

**Table S6: Primers for Enhancer cloning and gene expression and SDM primers for mutagenesis.**

| Enhancer name | Forward primer | Reverse primer |
|---|---|---|
| Sox2 (SRR111.1)* | GTGGGGTACAGGACATTGAAA | CTAGTCACTCATCCCCATGC |
| Med13l | CTCATCTGTGGTGTCGTTGTTG | GTTCAGGAGGGAGAGTTCTGTG |
| Sall1_MTL52 | TAAGCTGTGATGGCCTTATTGC | GAACCATTCTCATCTGACTCTGC |
| Sall1_MTL52core | GGTGTGCAGTGGGGATGGGGT | TGCCTGGGTAAGAAGGTCCC |
| Sall1_MTL40 | GAGTCTCTTCAGGACAACACCAT | GTCATCTAACAGCAAGCGAATCC |
| Sall1_MTL28 | GGCACCTCTAGAAAGTAAGACCTG | AGCTACACAAAGGTGGGTATCATT |
| Sall1_C1 | CTGGGTGAAGTCATTTCTGAGAC | ATCAACATCCAGGTGTGTCTTCT |
| Sall1_C2 | AGTGAAAGGAGAACTGTTAGATGACC | CTTAGTAATAGGGGCAGCTTGG |
| Sall1_129 | CGTGGCCTTCTTGTCAATg | CAACAGTACTCTGAACTCCCCAgT |
| Sall1_Cast | CCGTGGCCTTCTTGTCAATa | CAACAGTACTCTGAACTCCCCAaT |

| Primer name | Sequence |
|---|---|
| ***Med13l* enhancer mutagenesis primers** | |
| Med13l_GBX2_F | agggcatctccttacgtaatggttattatggggggtgggaca |
| Med13l_GBX2_R | tgtcccacccccataataaccattacgtaaggagatgccct |
| Med13l_SP1_1_F | agggtcatctgagaggagttaacctcaggtagaaaacatc |
| Med13l_SP1_1_R | gatgttttctacctgaggttaactcctctcagatgaccct |
| Med13l_ZIC3_1_F | ggtgacagtcagcatcaaacactgtgcaggttgggt |
| Med13l_ZIC3_1_R | acccaacctgcacagtgtttgatgctgactgtcacc |
| Med13l_TEAD4_F | gtggactccacctggcagtacggatttacagaatggg |
| Med13l_TEAD4_R | cccattctgtaaatccgtactgccaggtggagtccac |
| Med13l_FOXH1_F | ccctcctgtgtagtggggcctccacctggcattc |
| Med13l_FOXH1_R | gaatgccaggtggaggccccactacacaggaggg |
| Med13l_TFAP2C_F | ccacttccctggattcagtcccctaccggtg |
| Med13l_TFAP2C_R | caccggtaggggactgaatccagggaagtgg |
| Med13l_ZIC3_2_F | aaaatgtaaatctacccaatgcagcccccatacctcttgg |
| Med13l_ZIC3_2_R | ccaagaggtatgggggctgcattgggtagatttacatttt |
| Med13l_Pou5f1::Sox2_F | cttcctcgggcttttaaaagataaatctaccccctgcagc |
| Med13l_Pou5f1::Sox2_R | gctgcagggggtagatttatcttttaaaagcccgaggaag |
| Med13l_SP1_2_F | ccgaggctagagcatactccccacctgcc |
| Med13l_SP1_2_R | ggcaggtggggagtatgctctagcctcgg |
| ***Sox2* enhancer mutagenesis primers** | |
| Sox2_SRR111_NFIL3_F | gaatccgaggccttagcgccgaaacaggttcgagac |
| Sox2_SRR111_NFIL3_R | gtctcgaacctgtttcggcgctaaggcctcggattc |
| Sox2_SRR111_SP1_1_F | gcactcagggggcttatgcaggagcatcagg |
| Sox2_SRR111_SP1_1_R | cctgatgctcctgcataagccccctgagtgc |
| Sox2_SRR111_SP1_2_F | cttccggtaggggtgtatcggagggactgcaac |
| Sox2_SRR111_SP1_2_R | gttgcagtccctccgatacacccctaccggaag |
| Sox2_SRR111_TFAP2C_1_F | ccaaaccaagcacagcaccaatgtagtcagctaggtct |

| | |
|---|---|
| Sox2_SRR111_TFAP2C_1_R | agacctagctgactacattggtgctgtgcttggtttgg |
| Sox2_SRR111_GBX2_F | caggttctttttttaaaccctacgtgtcctccaccttccatttgag |
| Sox2_SRR111_GBX2_R | ctcaaatggaaggtggaggacacgtagggtttaaaaaaagaacctg |
| Sox2_SRR111_ZNF263_1_F | ttttaaaccctaattgtcctccacgtaccatttgagtcattcttaaattct |
| Sox2_SRR111_ZNF263_1_R | agaatttaagaatgactcaaatggtacgtggaggacaattagggtttaaaa |
| Sox2_SRR111_STAT1_F | ggcccatcccaggtgatttttttaaaccctaattgtcctccac |
| Sox2_SRR111_STAT1_R | gtggaggacaattagggtttaaaaaaatcacctgggatgggcc |
| Sox2_SRR111_TFAP2C_2_F | ccttcctctaggcagatccagtgggtttacaactggc |
| Sox2_SRR111_TFAP2C_2_R | gccagttgtaaacccactggatctgcctagaggaagg |
| Sox2_SRR111_ZNF263_2_F | tctcctccagctccctgctctaggcagctcc |
| Sox2_SRR111_ZNF263_2_R | ggagctgcctagagcagggagctggaggaga |
| Sox2_SRR111_SOX3_F | gacctccccctgtggtttctaagctctcctccagct |
| Sox2_SRR111_ SOX3_R | agctggaggagagcttagaaaccacagggggaggt |
| Sox2_SRR111_OCT4:SOX2_F | gagtggggtacaggacactgaagttgagagagggtttcttgtaa |
| Sox2_SRR111_OCT4:SOX2_R | ttacaagaaaccctctctcaacttcagtgtcctgtaccccactc |
| Sox2_SRR111_ESRRB_F | caccaggttatctctggcttcatgatgaccctgact |
| Sox2_SRR111_ESRRB_R | ccagtcagggtcatcatgaagccagagataacctggtg |
| ***Sall1* MTL 52 mutagenesis primers** | |
| +OS_F | CCCATTTGCATAACAAAGCAGGAGATTTGTTAC |
| +OS_R | CCTGCTTTGTTATGCAAATGGGGGGAGGGTG |
| +ETS_F | CAATGAACGTGTTGCTTTGGGTGGTGCTACTATGACCTTGAGGTCCCCACAGG |
| +ETS_R | ACCCAAAGCAACACGTTCATTGTTTCTGCCTGCCCCGTCTAGACACCCAATGCAAACATGACTGGTTTGAACTGG GCTTTCGGGTTGGG |
| +ETS_-FOXP1_F | caaaattaatacgaggcaaatttttttgtggttaaacacaatgtcttctacaccaggag |
| +ETS_-FOXP1_R | ctcctggtgtagaagacattgtgtttaaccacaaaaaaaatttgcctcgtattaattttg |
| +TS_F | cctgtggggacctccattccatagtagcaccacc |
| +TS_R | ggtggtgctactatggaatggaggtccccacagg |
| +TS_+E2F1_F | cacacatcagcagtaacaaacctcccgccctgttctgggaatggggggag |
| +TS_+E2F1_R | Ctccccccattcccagaacagggcgggaggtttgttactgctgatgtgtg |

* Only the 5’ half of 111 was used which corresponded to the TF bound region.

**Table S7: Guide RNAs.**

| Deletion | Sequence |
|---|---|
| Sall1_ΔEC | 5’ gRNA GTCTAGTGGT CTCTATAGCG (tgg)<br>3’ gRNA GCAGAGCTAC ACCCTCCGGA G (ggg) |
| Sall1_Δ1 | 5’ gRNA GTCTAGTGGT CTCTATAGCG (tgg)<br>3’ gRNA GGGTGAGCGA ACGAGCTTGG (agg) |
| Sall1_Δ2 | 5’ gRNA GGGTGAGCGA ACGAGCTTGG (agg)<br>3’ gRNA CTTTTCTTGG AGACCGGGCG (ggg) |
| Sall1_Δ3 | 5’ gRNA CTTTTCTTGG AGACCGGGCG (ggg)<br>3’ gRNA GCAGAGCTAC ACCCTCCGGA G (ggg) |
| Sall1_Δ2-3 | 5’ gRNA GGGTGAGCGA ACGAGCTTGG (agg) |

| | |
|---|---|
| | 3’ gRNA GCAGAGCTAC ACCCTCCGGA G (ggg) |
